# Alternative super-enhancers result in similar gene expression in different tissues

**DOI:** 10.1101/329987

**Authors:** Dóra Bojcsuk, Gergely Nagy, Bálint László Bálint

**Author notes:** These authors contributed equally to this work (Dóra Bojcsuk, Gergely Nagy).

## Abstract

Super-enhancers (SEs) are clusters of highly active enhancers, regulating cell type-specific and disease-related genes, including oncogenes^1–3^. The individual regulatory regions within SEs might be simultaneously bound by different transcription factors (TFs) and co-regulators such as P300, BRD4 and Mediator, which together establish a chromatin environment conducting to effective gene induction^4–6^. While cells with distinct TF profiles can have different functions, an unanswered question is how different cells control overlapping genetic programmes. Here, we show that the construction of oestrogen receptor alpha (ERα)-driven SEs is tissue specific, and both the collaborating TFs and the active SE components are largely differing between human breast cancer-derived MCF-7 and endometrial cancer-derived Ishikawa cells; nonetheless, SEs common to both cell types have similar transcriptional outputs. In the MCF-7 cell line, ERα-dominated SEs are also driven by the well-known FoxA1 and AP2γ TFs, as described previously^7^, whereas in Ishikawa cells, FoxM1, TCF12 and TEAD4 are as important as ERα for SE formation. Our results show that SEs can be constructed in several ways, but the overall activity of common SEs is the same between cells with a common master regulator. These findings may reshape our current understanding of how these regulatory units can fine-tune cell functions. From a broader perspective, we show that systems assembled from different components can perform similar tasks if a common functional trigger drives their assembly.

## Text

Understanding the structure of SEs is indispensable to understanding the regulation of targeted genes that most likely control cell function^3^. Although several regulatory factors contribute to the formation of SEs^8–10^, the activation of a dominant TF and its DNA-binding elements largely determine this process^7^. Previous studies focused mostly on temporal changes of SEs during differentiation but not on the transcriptional output of the active components of SEs common in different cell types^11–15^. The MCF-7 and Ishikawa female cancer cell lines derive from two different tissues but share ERα as the most important TF that regulates transcriptional processes upon hormonal stimulation (17β-oestradiol, E2). Therefore, we chose these two cancer models to compare their steady-state SE components, and processed publicly available ChIP-seq (chromatin immunoprecipitation coupled with sequencing) data to investigate how distinct genetic programmes are performed based on their cell line-specific, ERα-driven SEs.

We first assessed the ERα binding and the enriched motifs at the most active regions specific for MCF-7 and Ishikawa cells. Although both cell types had tens of thousands of ERα transcription factor binding sites (TFBSs), most of these binding sites, including the SE constituents, were characteristic of only one investigated cell line **(Fig. 1a, b and Supplementary Fig. 1a-d)**. The cell line-specific, ERα-driven SE constituents were ∼3.4-times more abundant in MCF-7 (n = 3,872) and ∼1.9-times more abundant in Ishikawa (n = 2,138) cells than constituents that were present in both cell lines (n = 1,124) **(Fig. 1b and Supplementary Fig. 1c, d)**. The presence of active chromatin (DNase I hypersensitivity), histone (H3K27ac) and enhancer (P300) marks followed these well-separated binding patterns (“clusters”), indicating that common and cell type-specific enhancers are indeed located within open and active chromatin regions **(Supplementary Fig. 1e, f)**. The first difference observed between the three clusters was seen in their enriched DNA motifs **(Fig. 1c and Supplementary Fig. 1g)**. Within the commonly occupied TFBSs, only the oestrogen response element (ERE) and different direct repeats (DRs) of the nuclear receptor (NR) half site were enriched, whereas in the cell type-specific clusters the motifs of other TFs were also enriched. Specifically, Fox and AP2 motifs were enriched in the MCF-7-specific cluster, and TEAD, TCF, AP-1 and SIX motifs in the Ishikawa-specific cluster, which did not show enrichment of the ERE motif but only the more general NR half site. FoxA1 plays a pioneering role in ERα function and AP2γ stabilizes ERα binding in breast cancer cells^16–18^, TEAD4 and TCF12 are coregulators of ERα in endometrial cancer cells^19^. Cooperation between TEAD4 and AP-1 has been reported in relation to transcriptional processes during tumourigenesis^20^; moreover, increased expression of SIX1 is a biomarker in human endometrial cancers^21^. These observations suggest a different mode of action between the SEs of our two chosen models.

**Figure 1.**
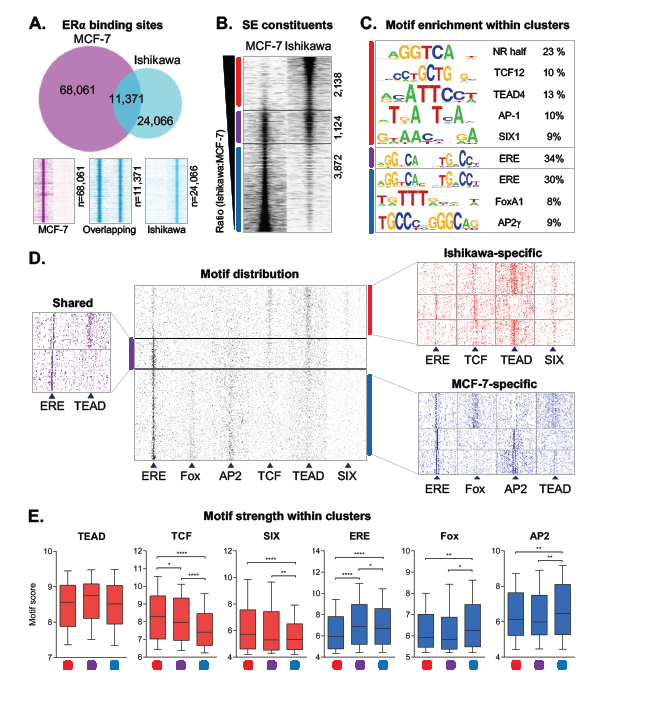
ERα-driven super-enhancer constituents show distinct binding patterns and motif preferences in MCF-7 and Ishikawa cells. **a,** Upper panel: an area-proportional Venn diagram showing the overlap between all ERα TFBSs upon E2 treatment in MCF-7 and Ishikawa cells. Lower panel: read distribution plots showing ERα binding at the shared and cell line-specific TFBSs upon E2 treatment in 2-kb frames. **b,** A read distribution plot showing ERα density on ERα-driven super-enhancer (SE) constituents derived from MCF-7 and Ishikawa cells in 2-kb frames. Peaks were sorted based on the ratio of RPKM (reads per kilobase per million mapped reads) values calculated from Ishikawa and MCF-7 cells and were separated into three different clusters: the red line represents Ishikawa-specific constituents (n = 2,138), the purple line represents shared constituents (n = 1,124) and the blue line represents MCF-7-specific SE constituents (n = 3,872). **c,** The enriched motifs and their target percentages within the three clusters. **d,** The motif distribution plot of ERE, Fox, AP2, TCF, TEAD and SIX motifs in 1.5-kb frames around the summit position of ERα-driven SE constituents in the same order as introduced in Figure 1b (middle). Coloured heat maps represent shared and cell line-specific clusters when peaks were further clustered based on the presence or absence of the most frequent motifs. **e,** Box plots showing the distribution of motif strengths within the three main clusters introduced in Figure 1b. The boxes represent the first and third quartiles, the horizontal lines indicate the median scores and the whiskers indicate the 10^th^ to 90^th^ percentile ranges. Paired t-test, * significant at *P* < 0.05, ** at *P* < 0.01, *** at *P* < 0.001, **** at *P* <0.0001.

Based on these initial findings, we carried out a detailed investigation of how different TFs contribute to the formation of both cell type-specific and shared ERα-driven SEs. First, as a validation, we mapped the matrix of identified DNA motifs and found that the shared ERα binding sites showed large numbers of EREs and smaller numbers of TEAD, TCF and SIX elements, whereas the cell type-specific enhancers showed expected motif distribution patterns: Fox and AP2 motifs were enriched at the MCF-7-specific, and TCF, TEAD and SIX motifs were enriched at the Ishikawa-specific ERα binding sites **(Fig. 1d, Supplementary Fig. 2a)**. The creation of sub-clusters based on motif distribution showed that certain motifs (e.g., ERE and TEAD or Fox and AP2) might mutually exclude each other **(Fig. 1d)**. To further examine the motif specificity of cell type-specific binding sites, we plotted the motif strengths within the cell type-specific and shared clusters **(Fig. 1e, Supplementary Fig. 2b)**. Generally, the motif strengths correlated well with the motif distribution patterns; however, the top TEAD motifs were within the shared cluster and not at the Ishikawa-specific sites. The above analyses pointed out that the two cell types use different sets of TFs, but even a common TF might show distinct binding pattern.

**Figure 2.**
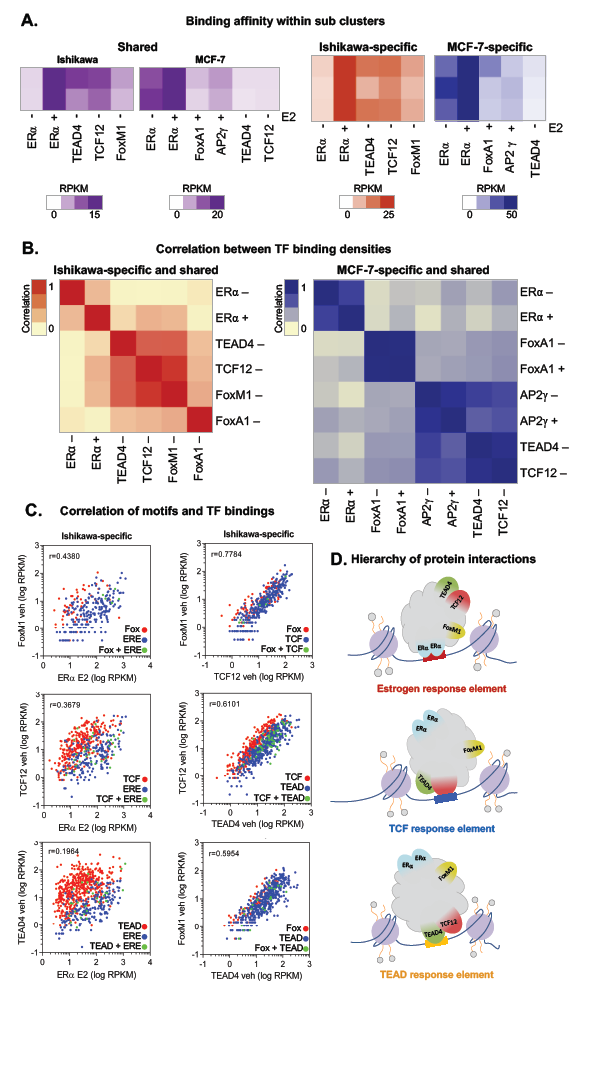
Response elements determine a consistent hierarchy between transcription factor binding events. **a,** Heat maps showing the density of relevant TFs in the presence (+) or absence (-) of E2 within the same sub-clusters introduced in Figure 1d. The plotted densities are the averages of the values calculated by Homer within 50-bp regions around the summit of the sub-clusters’ SE constituents. In the case of shared (common) peaks, ChIP-seq coverages were separately calculated for both Ishikawa and MCF-7 cells. **b,** Correlation plots showing the correlation coefficients (r) calculated from the densities of all investigated TFs on the SE constituents (summit ± 50-bp regions) of Ishikawa and MCF-7 cells. **c,** Scatter plots showing the densities of the indicated TFs (upon vehicle [veh] or E2 treatment) on their DNA-binding motifs within the MCF-7- or Ishikawa-specific ERα-driven SE constituents. Red and blue dots represent protein binding on a specific single motif, and green dots represent protein binding on a region with the motifs of both examined TFs. **d,** Working models of the supposed hierarchy between ERα, FoxM1, TCF12 and TEAD4 TFs in Ishikawa cells based on the presence of ERE, TCF or TEAD response elements.

TF motifs can be bound by several proteins of a TF family; therefore, we compared the expression levels of all members of the emerging TF families from publicly available RNA-seq data sets **(Supplementary Fig. 1e, 3a)**. Not only *FOXA1* and *TFAP2C* (encoding AP2γ) but also *ESR1* (encoding ERα) showed much lower expression in Ishikawa cells than in MCF-7 cells. Out of the more than 40 members of the Fox family, *FOXM1* had the highest expression in Ishikawa cells; however, *FOXD1* also showed a notable expression level **(Supplementary Fig. 3a, b)**. *TCF12* and *TEAD4* showed higher expression in Ishikawa cells than in MCF-7 cells, but this was also true for other family members, such as *TCF3* and *TEAD2*. *SIX* genes were lowly expressed in both cell lines and in this comparison, *SIX5* rather expressed in Ishikawa cells and *SIX4* was rather specific to MCF-7 cells. The performed gene expression comparison confirmed the role of the collaborating TFs and above these, highlighted FoxM1 as a TF with major role in Ishikawa cells.

**Figure 3.**
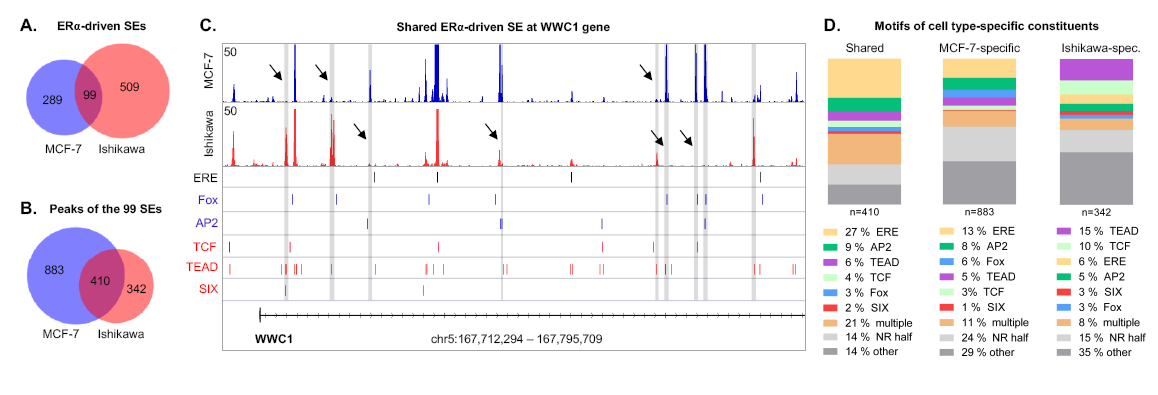
Shared ERα-driven super-enhancers are composed of different transcription factor binding sites in MCF-7 and Ishikawa cells. **a, b,** Area-proportional Venn diagrams showing the overlap between all ERα-driven SEs of MCF-7 and Ishikawa cells **(a)** and the overlap between the constituents of the 99 shared SEs **(b)**. **c,** The Integrative Genomics Viewer snapshot of ERα ChIP- seq coverage on the WWC1 locus showing an SE that is formed upon E2 treatment in both MCF-7 and Ishikawa cells (top). The interval scale is 50. The matrix of ERE, Fox, AP2, TCF, TEAD and SIX motifs was mapped within the summit ± 50-bp regions of the ERα peaks, and the indicated putative elements are represented as thin lines (bottom). Peaks marked with arrows and highlighted in grey show different binding patterns between MCF-7 and Ishikawa cells. **d,** The proportion of investigated DNA motifs within the 99 shared ERα-driven SEs visualized on three bar charts (stacked up to 100%) and classified according to Figure 3b.

The overall TF binding densities generally followed the motif distribution-based sub-clusters defined in Figure 1b and d **(Supplementary Fig. 3c)**. Recruitment of ERα upon E2 treatment was seen in each sub-cluster, even in binding sites that lacked ERE **(Fig. 2a, Supplementary Fig. 1e)**. To examine the protein-protein interactions suggested by these results, we performed a correlation analysis on TF binding **(Fig. 2b)**. In Ishikawa cells, FoxM1 and TCF12 showed the strongest co-occurrence both with each other and with TEAD4 and E2-induced ERα. The correlation heat map for MCF-7 suggests the independent binding of key TFs. To examine both the protein-protein and DNA-protein concomitance, we plotted TF densities at their putative TFBSs **(Fig. 2c and Supplementary Fig. 3d)**. This kind of visualization clearly demonstrated that different TFs show higher density at their own elements, but at the same time, we obtained information about their “affinity” to each other. In Ishikawa cells, the presence of ERα correlated best with FoxM1 binding, followed by binding with TCF12 and TEAD4, and pairwise comparisons also suggested a FoxM1/TCF12, TCF12/TEAD4 and FoxM1/TEAD4 interaction, which implies a tripartite complex interacting with ERα **(Fig. 2c, d and Supplementary Fig. 3d)**. A contact between a steroid hormone receptor and a Fox protein is not unprecedented as androgen receptor (AR) and FoxA1 associate to form an ARE::Fox composite element; however, usually a single ARE or Fox motif is sufficient for binding by both proteins^22–24^. In our proposed mechanism, any TF can bind its DNA element, although the Fox motifs are very rare. The TCF12/TEAD4 relationship was also reproduced with lower protein levels in MCF-7 cells **(Supplementary Fig. 3d)**. This means that in Ishikawa cells, there is no need for direct DNA binding by ERα for regulation (as we described previously in MCF-7 cells). Instead, certain TF partners can make ERα a hormone-sensitive coregulator, which process also increases TF-binding affinity upon ligand treatment **(Fig. 2a, b)**. In MCF-7 cells, there was no tight co-occupancy between dominant proteins, but ERα/FoxA1 concomitances seemed to be the least frequent. Upon E2 treatment, we observed a slight recruitment of FoxA1 and a stronger recruitment of AP2γ, as has been described previously **(Supplementary Fig. 3d)**. There were few TFBSs where two motifs could be mapped (green dots); these regions were usually bound by their specific TFs to a similar extent. Together, these findings indicate that the TF binding is follows well the DNA motif pattern and there is a well-defined cooperativity and hierarchy between the TFs promoting the formation of complexes on SEs.

By focusing on entire SE regions, we found 99 SEs that partly or fully overlapped between MCF-7 and Ishikawa cells, but these “common” SEs shared only a quarter of their ERα TFBSs (410 in total) **(Fig. 3a-c and Supplementary Fig. 4a)**. These commonly used (shared) binding sites were dominated by ERE alone (27%) or in combination with other motifs (21%, multiple) and NR half sites (14%) **(Fig. 3d and Supplementary Fig. 4b)**. The MCF-7-specific binding sites showed a similar motif distribution but with a considerably higher proportion of NR half sites (24%) and other motifs (29%), whereas in Ishikawa cells, TEAD (15%), TCF (10%) and NR half motifs (15%) dominated compared to ERE **(Fig. 3d and Supplementary Fig. 4b)**. Ishikawa-specific SEs were typically bound by ERα at EREs not only in Ishikawa but also in MCF-7 cells, whereas MCF-7-specific SEs were rarely bound in Ishikawa cells **(Supplementary Fig. 4c, d)**. This result is consistent with the notion that, while in MCF-7 cells ERα is dominantly recruited to EREs or NR half sites, in Ishikawa cells ERα can also be recruited by TEAD4 and/or TCF12 **(Fig. 3d)**.

**Figure 4.**
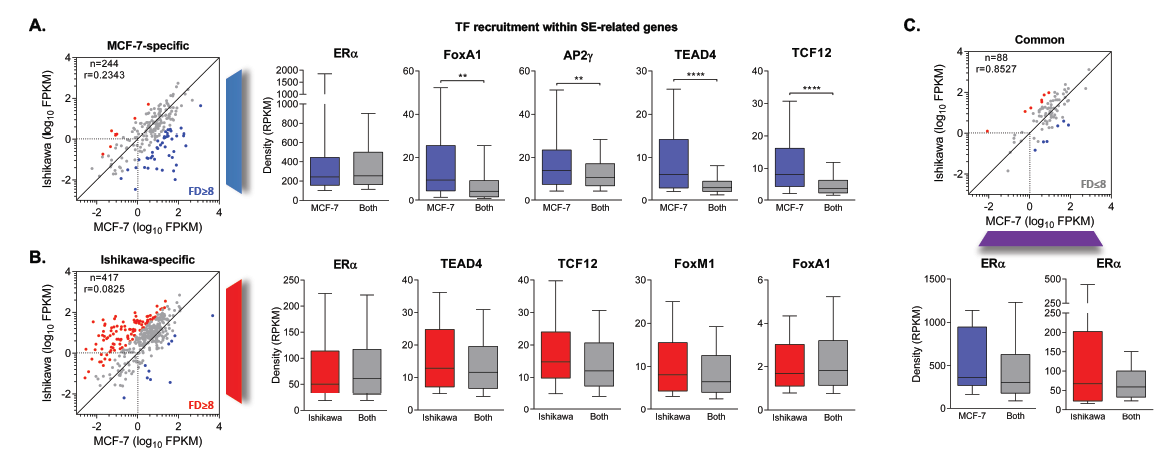
Genes regulated by shared SEs show identical expression in MCF-7 and Ishikawa cells. Scatter plots showing the expression levels of genes closest to the MCF-7-specific **(a)**, Ishikawa-specific **(b)** and shared **(c)** ERα-driven SEs. Grey dots represent genes within an eight-fold difference (FD) range; blue and red dots represent genes that exceed this range and are specific to MCF-7 or Ishikawa cells, respectively. The box plots show the average densities of the indicated TFs within SEs related to the differentially (blue or red boxes, FD ≥ 8) or similarly (grey boxes, FD ≤ 8) regulated genes. The boxes represent the first and third quartiles, the horizontal lines indicate the median coverage values and the whiskers indicate the 10^th^ to 90^th^ percentile ranges. Paired t-test, * significant at *P* <0.05, ** at *P* < 0.01, *** at *P* < 0.001, **** at *P* < 0.0001. Coverage (RPKM, reads per kilobase per million mapped reads) values were calculated within 100-bp regions around the summit of the ERα peaks.

In the last step, we compared the expression levels of the genes regulated by cell type-specific and shared SEs in the two cell lines. Surprisingly, only a fraction of the MCF-7-specific SE regulated genes showed considerably higher (fold difference ≥ 8) expression in MCF-7 cells than in Ishikawa cells **(Fig. 4a)**. A similar phenomenon was observed for the Ishikawa-specific SE related genes **(Fig. 4b)**. To dissect the contribution of individual TFs to gene expression profiles, we further investigated the protein densities of the polarizing (blue or red) and less deterministic SEs **(Fig. 4a-c)**. SEs of MCF-7-specific genes were covered by significantly more TF, except for ERα, than those associated with genes expressed at a similar level in both cell types **(Fig. 4a)**. We detected similar tendencies for the regulatory regions of Ishikawa-specific genes, although these differences were not significant **(Fig. 4b)**. Genes regulated by the shared SEs showed similarly high expression and generated similar transcriptional output in the MCF-7 and Ishikawa cell lines, even though they had largely different sets of collaborative factors **(Fig. 4c)**. These results suggest that, although ERα is highly enriched at the SEs of target genes, their gene expression level can be further improved by its collaborating factors.

Finally, we were curious whether the basic findings regarding SE constituents and their DNA motifs can be observed in primary tumour cells of patients diagnosed with different stages of breast cancer. We found that most of the ERα-driven SEs constituents of a tamoxifen-responder, a non-responder and a metastatic patient are clustered separately; however, common peaks can be seen regardless of whether they are constituents of a SE or not in the other tumour type **(Supplementary Fig. 5a, b)**. By using each SE peaks (11,385 in total), near the ERE, motifs of Fox, AP-1 and NF-1 TFs are enriched **(Supplementary Fig. 5d)**. Mapping them together with the previously identified breast cancer-specific AP2 motif, the result reflected patient- or stage-specific signatures: while motifs of AP2, AP-1 and the neurofibromin NF-1^25^, which can modulate the response to tamoxifen, are not highly enriched in metastasis, Fox motif is characteristic of it **(Supplementary Fig. 5e, f)**. These findings prove that ERα-driven SEs have a patient- or stage-specific motif composition, therefore, the regulatory layer of the genomic code is interpreted in different ways in patient-derived samples, as well.

In conclusion, our study highlights the differences in the role of ERα between the SEs of two E2-sensitive cell lines and in primary breast cancer samples: while in MCF-7 cells, ERα has no coequal TF partners **(Fig. 1c, d and 3d)**, in Ishikawa cells, we can assume the existence of at least a tripartite protein complex composed of TEAD4, TCF12 and FoxM1 in which FoxM1 might have the highest affinity to ERα **(Fig. 2b-d)**. Importantly, ERα does not seem to be an activator itself as the density of only its collaborating TFs correlates with gene expression, independent of TF classes **(Fig. 4)**. Our results also suggest that SEs are dynamic structures and that different tissues or even cancer subtypes can assemble them in different ways. This novel layer of genomic regulation encoded in the DNA sequence itself must be considered to understand how our genome works and how these codes are translated by SEs into tissue-specific genetic programs.

## Methods

### Data selection

Raw ChIP-seq, DNase-seq and RNA-seq data were downloaded from the Gene Expression Omnibus (GEO). As we used data from ECC-1 isolates of ATCC (American Type Culture Collection, Manassas, VA) that were genotyped as Ishikawa cells^26^; we therefore referred to them as Ishikawa cells. Detailed information about the selected data (e.g., GEO identifiers and references) is included in Supplementary Fig. 1a, e and Supplementary Fig. 5a.

### ChIP-seq analysis

Raw sequence data were re-analyzed with an updated version of our previously published computational pipeline^27^ as follows: reads were aligned to the hg19 reference genome assembly (GRCh37) by using the Burrows-Wheeler Alignment (BWA) tool (v07.10)^28^, then BAM files were generated with SAMtools (v0.1.19)^29^. Coverage files were created by the *makeUCSCfile.pl* script of the Hypergeometric Optimization of Motif EnRichment (HOMER) package (v4.2)^30^, and peaks were predicted using the Model-based Analysis of ChIP-Seq (MACS2) tool (v2.0.10) with *-callpeak* parameter^31^. To remove the artifacts from the predicted peaks, we used the blacklisted genomic regions of the Encyclopedia of DNA Elements (ENCODE)^32^.

Reads Per Kilobase per Million mapped reads (RPKM) values were calculated on the ± 50-bp regions relative to the peak summits by using the *coverageBed* program of BedTools (v2.23.0)^33^. The number of overlapping peaks and regions was defined by using the DiffBind package (v1.2.4) in R^34^.

Read distribution (RD) heat maps were generated by *annotatePeaks.pl* with *-hist 50* and *-ghist* parameters (HOMER). Coverage values for average protein density heat maps were calculated on the summit positions of the RD plots.

### Super-enhancer prediction

Super-enhancers were predicted from the E2-treated ERα ChIP-seq samples applying the HOMER’s *findPeaks.pl* script and the *-style super* parameter. To generate “super-enhancer plot”, we used the *-superSlope -1000* parameter, and the thus generated “Normalized Tag Count” values were plotted. Tag counts (rpm/bp; reads per million per base pair) of the ERα (super-)enhancers were ranked by their ChIP-seq coverage. Definition of super-enhancers was based on the original strategy, where the outstandingly “active” enhancers or broader regions in which enhancers are closer than 12.5 kb to each other are over slope 1 in the rank order.

### Motif analysis

Motif enrichment analysis was carried out by the *findMotifsGenome.pl* script of HOMER. It was performed on the ±100 bp flanking regions of the peak summits. The search length of the motifs were 10, 12, 14 and 16 bp. *P*-values were calculated by comparing the enrichments within the target regions and that of a random set of regions (background) generated by HOMER.

For motif distribution plot, motif matrices (shown on Supplementary Fig. 2a) were mapped in 30-bp windows within 1.5-kb frame relative to the ERα peak summits using *annotatePeaks.pl* with *-mbed* parameter, BEDtools and other command line programs. Clustering of motif distribution patterns was done by Cluster 3.0. Top motif score for each examined regions were determined by *annotatePeaks.pl* with *-mscore* parameter. Thresholds of the plotted scores were selected before the last markedly high motif numbers.

### DNase-seq analysis

The primary analysis of DNase-seq data was carried out as described for ChIP-seq data.

### RNA-seq analysis

Raw sequence data were aligned to the hg19 reference genome assembly (GRCh37) by using TopHat (v2.0.7). The Fragments Per Kilobase of transcript per Million mapped reads (FPKM) values were calculated by Cufflinks (v2.0.2) with default parameters^35^.

### Gene annotation

Super-enhancers were annotated to the nearest transcription start site (TSS) of the protein coding genes by using PeakAnnotator^36^.

### Visualization

Read distribution, average protein density and correlation heat maps were plotted by Java TreeView (v1.1.6r4)^37^. Area-proportional Venn diagrams were produced by BioVenn^38^. Box plots, scatter plots, bar charts and histograms were created with GraphPad Prism 6. Coverage files were visualized by Integrative Genomics Viewer (IGV)^39^.

## Acknowledgement

This work was supported by University of Debrecen in the programme “Internal Research Grant of the Research University” entitled “Dissecting the genetic and epigenetic components of gene expression regulation in the context of the 1000 genomes project” and through the internal research funding provided by the Department of Biochemistry and Molecular Biology; B.L.B. is a Szodoray Fellow of the University of Debrecen, Faculty of Medicine and an alumni of the Magyary Zoltan fellowship supported by the TÁMOP 4.2.4.A/2-11-1-2012-0001 grant implemented through the New Hungary Development Plan co-financed by the European Social Fund and the European Regional Development Fund; D.B. is supported by the ÚNKP-17-3 New National Excellence Program of the Ministry of Human Capacities. G.N. is supported by the Hungarian Scientific Research Fund (OTKA) PD 124843. This study makes use of publicly available sequencing data, which were cited in the manuscript.

## Author Contributions

B.L.B., D.B. and G.N. designed the study. D.B. collected and analysed the data. D.B and G.N. carried out the detailed computational analysis and wrote the manuscript. B.L.B. revised the analysis and manuscript.

---

**Supplementary Figure 1.**
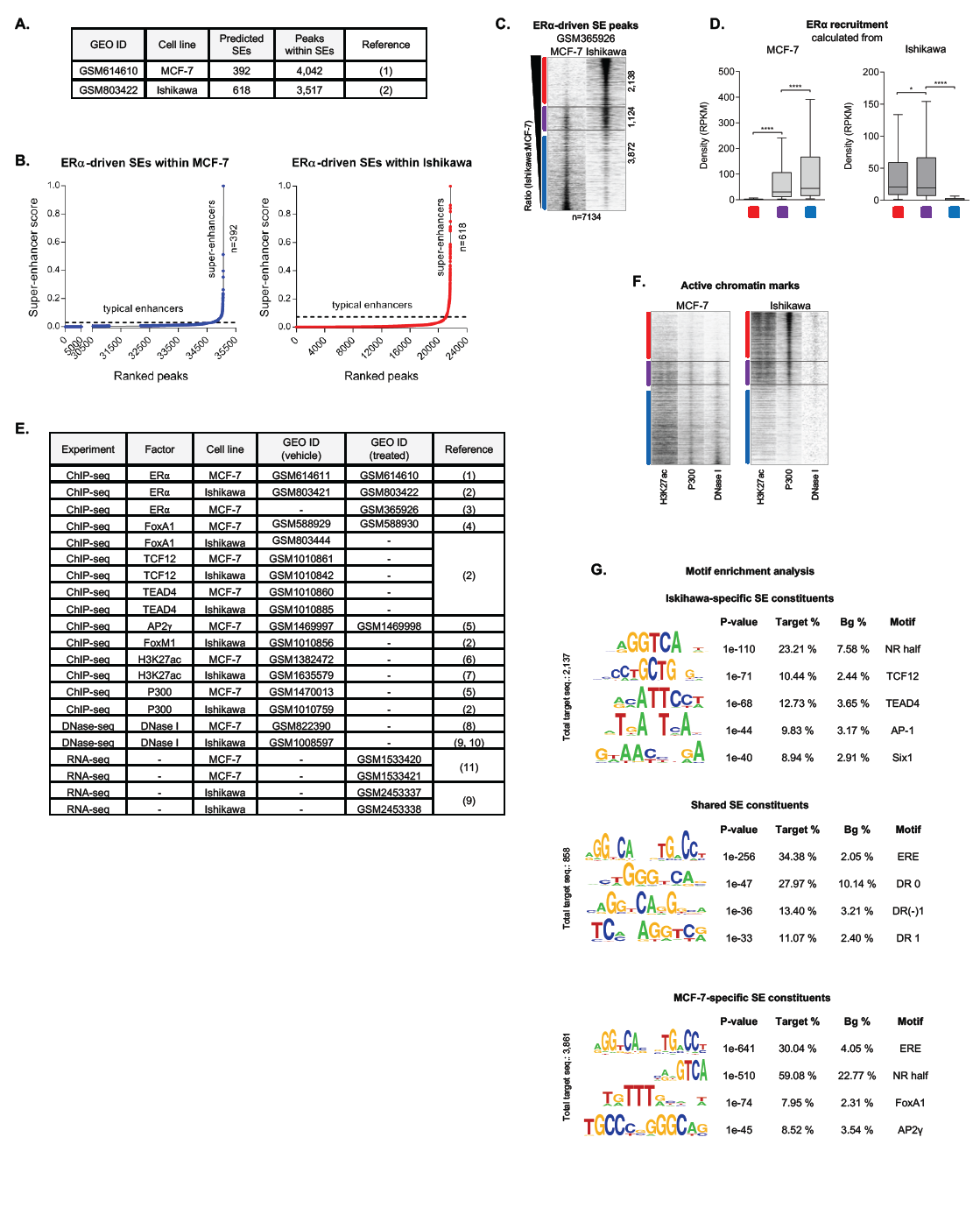
Enrichment of active chromatin marks and regulatory factors follows ERα binding patterns in MCF-7 and Ishikawa cells. **a,** Information about the ERα ChIP-seq samples used for the basic analysis. **b,** The definition of ERα-driven SEs in MCF-7 and Ishikawa cell lines. Groups of enhancers (or even single enhancers) over slope 1 were considered to be SEs. **c,** Read distribution plot showing the pooled peak set (n = 7,134) of ERα-driven SEs upon E2 treatment (GSM365926). Coverages were plotted in 2-kb frames. The order of peaks was determined from the GSM614610 (MCF-7) and GSM803422 (Ishikawa) data as introduced in Figure 1b. Despite the treatment conditions (GSM365926: 10 nM E2 for 1 h vs. GSM614610/GSM803422: 100 nM E2 for 45 min), the same tendencies can be observed. **d,** Box plots showing ERα recruitment within Ishikawa-specific, shared and MCF-7-specific clusters. RPKM (reads per kilobase per million mapped reads) values were calculated on the summit ± 50-bp regions of the ERα peaks, separately from the MCF-7 and Ishikawa ChIP-seq samples. The boxes represent the first and third quartiles, the horizontal lines indicate the median RPKM values and the whiskers indicate the 10^th^ to 90^th^ percentile ranges. Paired t-test, * significant at *P* < 0.05, ** at *P* < 0.01, *** at *P* < 0.001, **** at *P* < 0.0001. **e,** Information about ChIP-seq, DNase-seq and RNA-seq samples used for the characterization of ERα-driven SEs. **f,** Read distribution plots of H3K27ac and P300 ChIP-seq and DNase-seq (DNase I) data in MCF-7 and Ishikawa cell lines upon vehicle treatment relative to the ERα SE constituents in 2-kb frames in the same order as introduced in Figure 1b. **g,** Detailed motif enrichment results within the ERα peaks of the three clusters (related to Figure 1c). *P*-values and target and background (Bg) percentages are included for each motif.

**Supplementary Figure 2.**
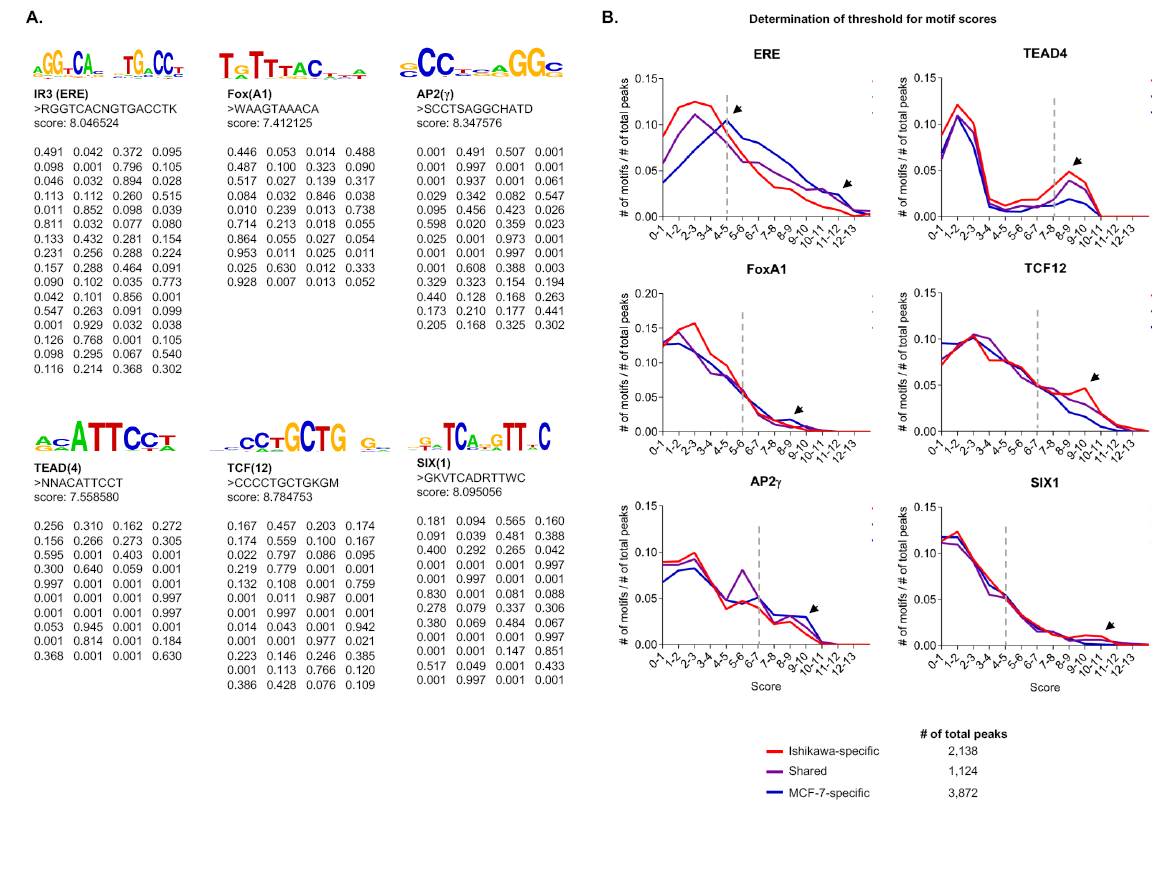
Transcription factor binding correlates well with response element strength. **a,** The logos and matrices of enriched ERE, Fox, AP2, TCF, TEAD and SIX motifs used for mapping. **b,** Histograms showing the frequency (#) of motifs depending on their score. The total number of motifs was divided with the given cluster size. Red, blue and purple lines represent Ishikawa-specific, MCF-7-specific and common ERα peaks, respectively. Dashed lines indicate the score threshold used for the motif strength analysis shown in Figure 1e, and arrows show motif enrichments specific to a cluster.

**Supplementary Figure 3.**
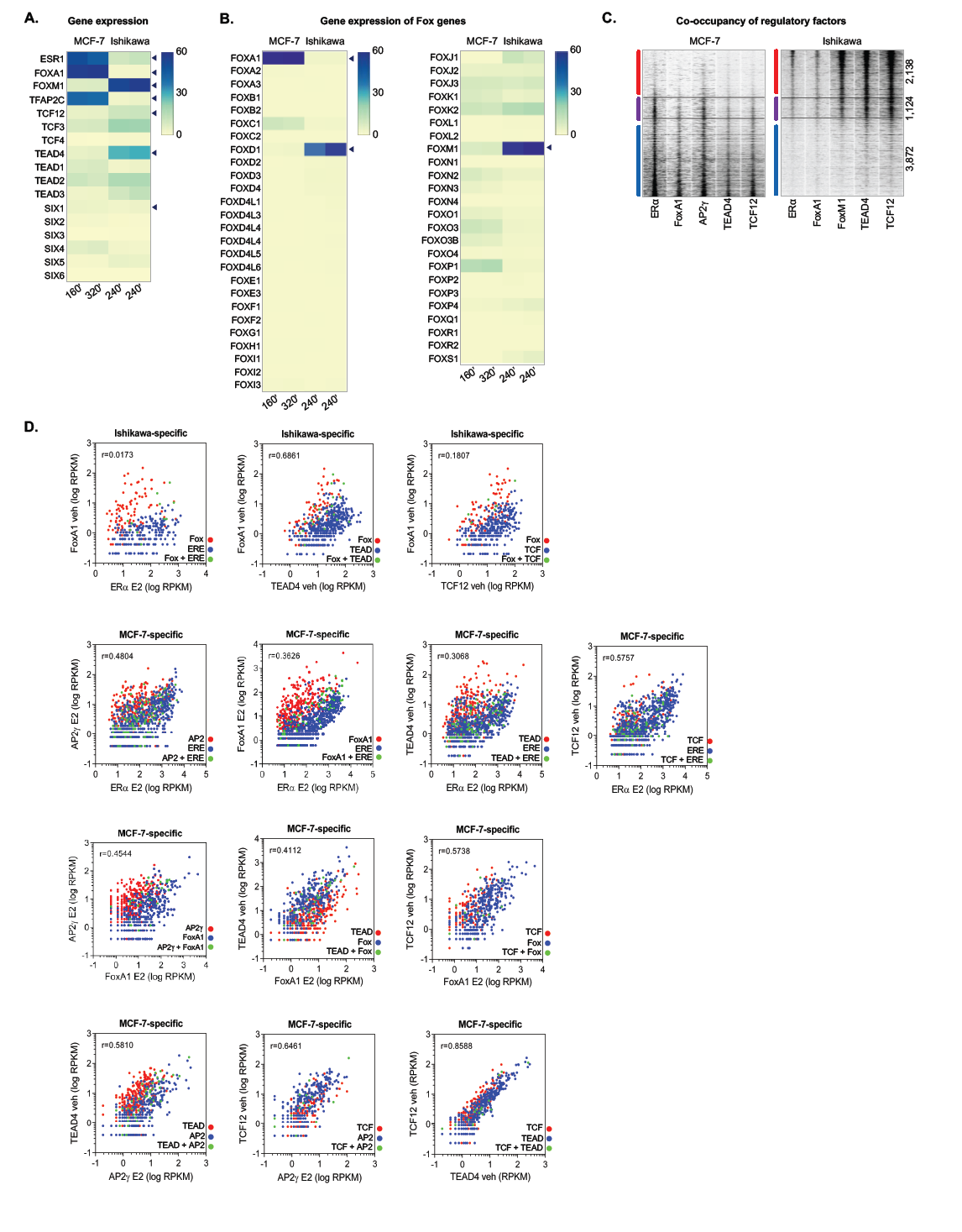
Discovering transcription factor interactions with their response elements. **a, b,** The gene expression levels of putative regulator TF families **(a)** and the whole Fox family **(b)** in MCF-7 and Ishikawa cells. MCF-7 cells were treated with 10 nM E2 for 160 or 320 min, and Ishikawa cells were treated with 10 nM E2 for 240 min. Fragments per kilobase per million mapped reads (FPKM) values are shown. **c,** Read distribution plots of the indicated TFs in MCF-7 and Ishikawa cells upon vehicle treatment in 2-kb frames on the regions introduced in Figure 1b. **d,** Scatter plots showing the densities of the indicated TFs (upon vehicle [veh] or E2 treatment) on their DNA-binding motifs within the MCF-7-or Ishikawa-specific ERα-driven SE constituents. Red and blue dots represent protein binding at the specific single motif, and green dots represent protein binding at a region with the motifs of both examined TFs.

**Supplementary Figure 4.**
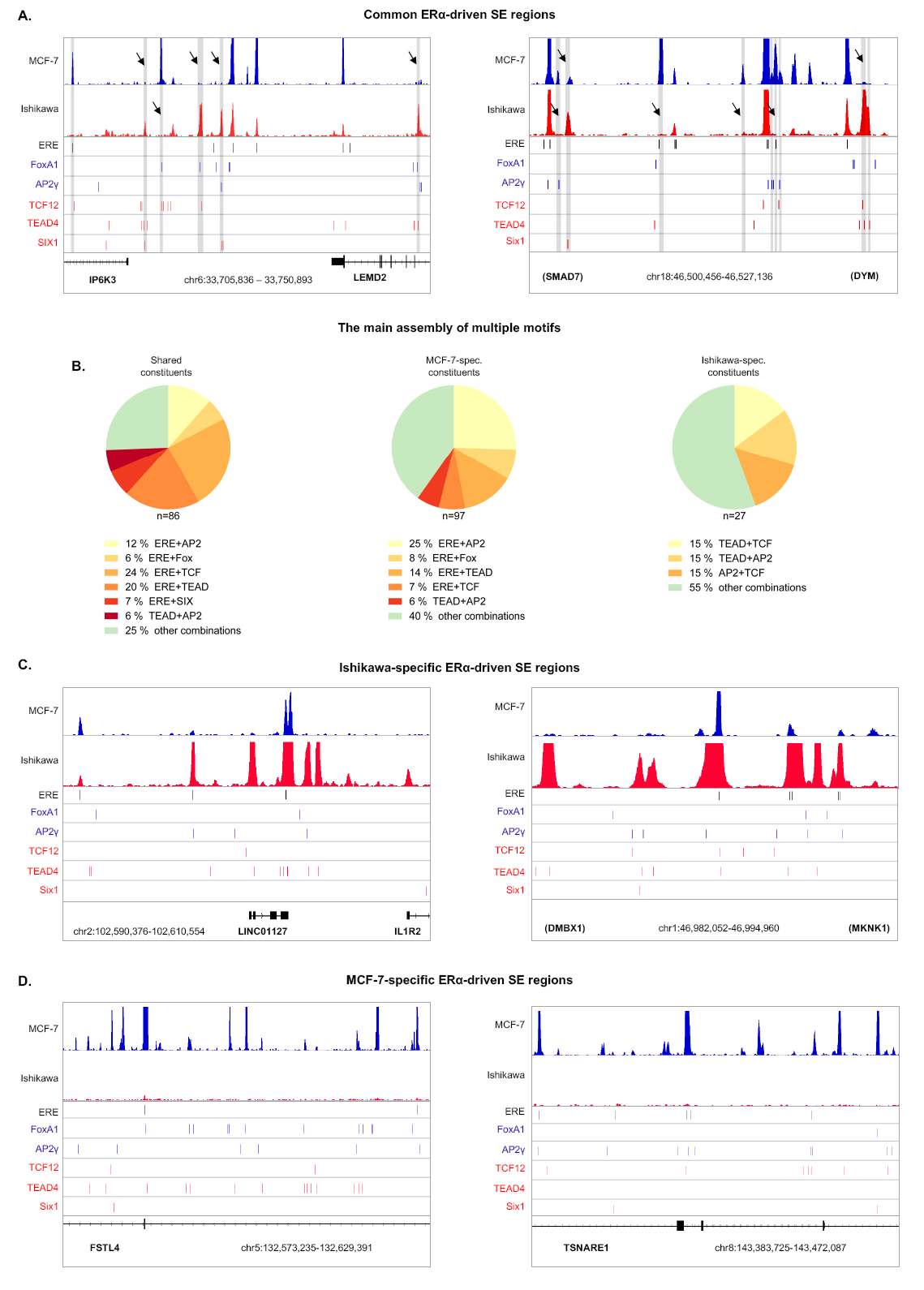
ERα-driven super-enhancers are driven by different motifs in MCF-7 and Ishikawa cells. **a, c, d** Integrative Genomics Viewer snapshots of ERα ChIP-seq coverage on overlapping (common) **(a)** Ishikawa-specific **(c)** and MCF-7-specific **(d)** ERα-driven SEs in MCF-7 and Ishikawa cells upon E2 treatment. The interval scale is 50. The matrix of ERE, Fox, AP2, TCF, TEAD and SIX motifs was mapped within the summit ± 50-bp regions of the ERα peaks, and the indicated putative elements are represented as thin lines (bottom). Peaks marked with arrows and highlighted in grey show different binding patterns between MCF-7 and Ishikawa cells. **b,** Frequency of the top multiple motif appearances within the shared, the MCF-7-specific and Ishikawa-specific constituents of overlapping SE regions.

**Supplementary Figure 5.**
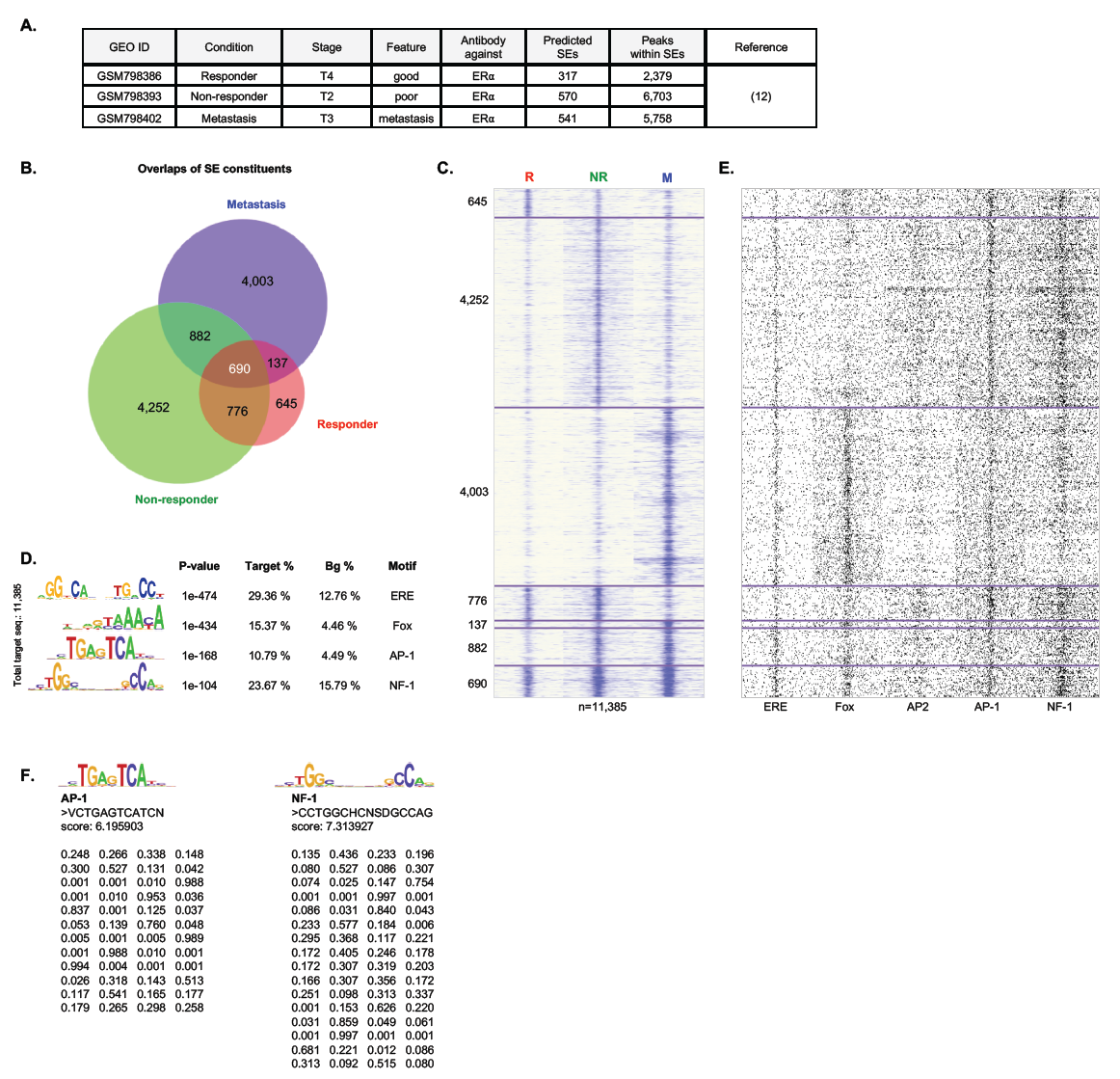
Different breast cancer stages are driven by distinct TFs and motifs. **a,** Information about the ERα ChIP-seq samples used for the analysis. **b,** Area-proportional Venn diagram showing the overlaps between the ERα-driven SE constituents of a tamoxifen-responder, a non-responder and a metastatic patient. **c,** Read distribution plot representing ERα density on stage-specific and overlapping ERα-driven SE constituents. Coverages were plotted in 2-kb frames. **d,** The enriched motifs within the entire set of tamoxifen-responder, non-responder and the metastatic SE peaks. *P*-values and target and background (Bg) percentages are included for each motif. **e,** The motif distribution plot of ERE, Fox, AP2, AP-1 and NF-1 motifs in 1.5-kb frames around the summit position of ERα-driven SE constituents in the same order as introduced in Supplementary Fig. c. **f,** The logos and matrices of newly enriched AP-1 and NF-1 motifs used for mapping. The mapped ERE, Fox and AP2 motif matrices were indicated on Supplementary Fig. 2a.

